## Supplementary Figures for "MSFragger-DDA+ Enhances Peptide Identification Sensitivity with Full Isolation Window Search"

| Tool | Target sequences | Entrapment sequences | FDP estimation | |
|---|---|---|---|---|
| | | | upper bound | lower bound |
| MaxQuant | 31146 | 54 | 0.35% | 0.17% |
| MetaMorpheus | 44319 | 689 | 3.06% | 1.53% |
| MSFragger-DDA FragPipe | 42726 | 22 | 0.10% | 0.05% |
| MSFragger-DDA+ FragPipe | 68629 | 45 | 0.13% | 0.07% |

**Supplementary Figure 1. Peptide-level FDP evaluation for MaxQuant, MetaMorpheus, MSFragger, and MSFragger-DDA+.** Two calculation methods, including the upper bound and lower bound method, were applied.

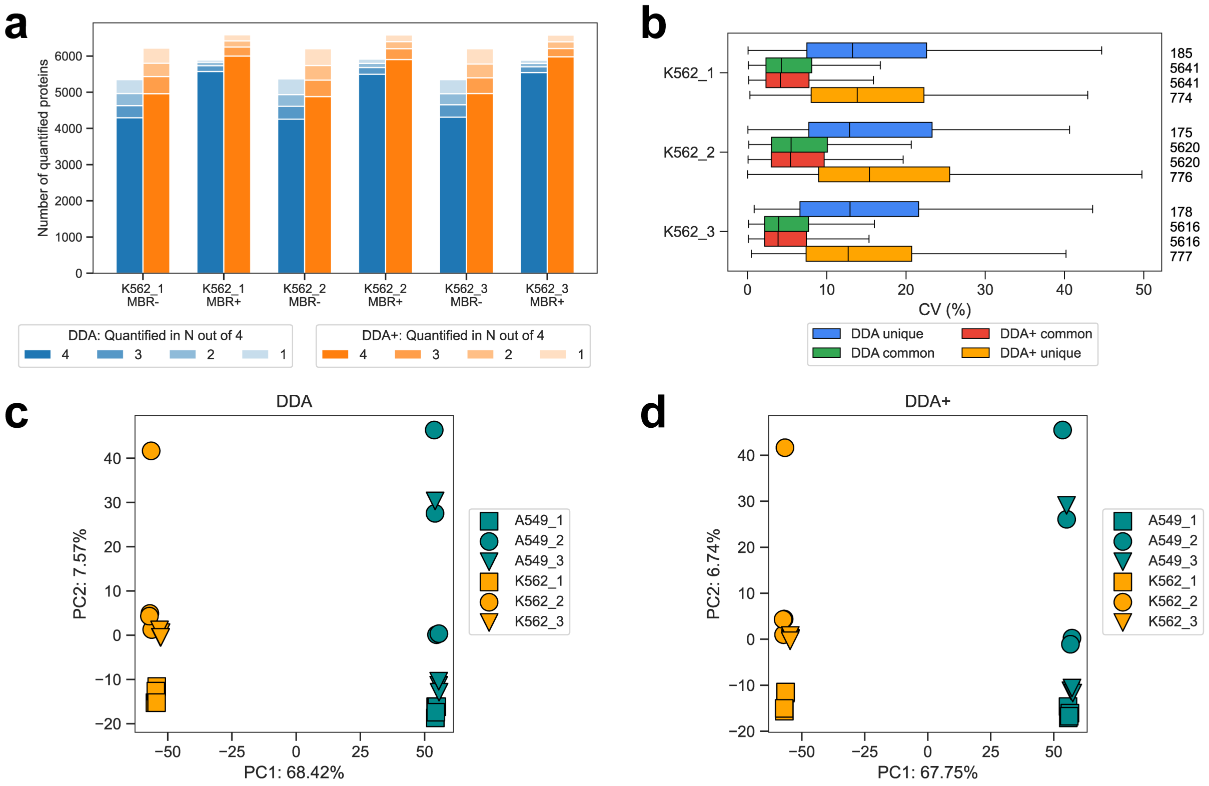


**Supplementary Figure 2. Performance benchmarking using timsTOF ddaPASEF data. (a)** Number of quantified proteins from the DDA and DDA+ workflows. The samples are from the K562 cell line. There are three biological replicates. Each biological replicate contains four technical replicates. “MBR+” and “MBR-” are with and without MBR, respectively. **(b)** Numbers and CVs of overlapped and non-overlapped proteins quantified from the DDA and DDA+ workflows using the K562 cell line. The blue box plots are from the unique proteins of the DDA mode, the green box plots are from the common proteins of the DDA mode, the red box plots are from the common proteins of the DDA+ mode, and the yellow box plots are from the unique proteins of the DDA+ mode. The common proteins are the overlapping proteins quantified in both DDA and DDA+ modes. The numbers on the right are the quantified proteins. The box in each plot captures the interquartile range (IQR) with the bottom and top edges representing the first (Q1) and third quartiles (Q3), respectively. The median (Q2) is indicated by a horizontal line within the box. The whiskers extend to the minima and maxima within 1.5 times the IQR below Q1 or above Q3. **(c)** PCA plot of the quantitative results from two cell lines, A549 and K562, and their three experimental replicates. The result is from the DDA workflow. **(d)** Similar to **(c)** but the result is from the DDA+ workflow.

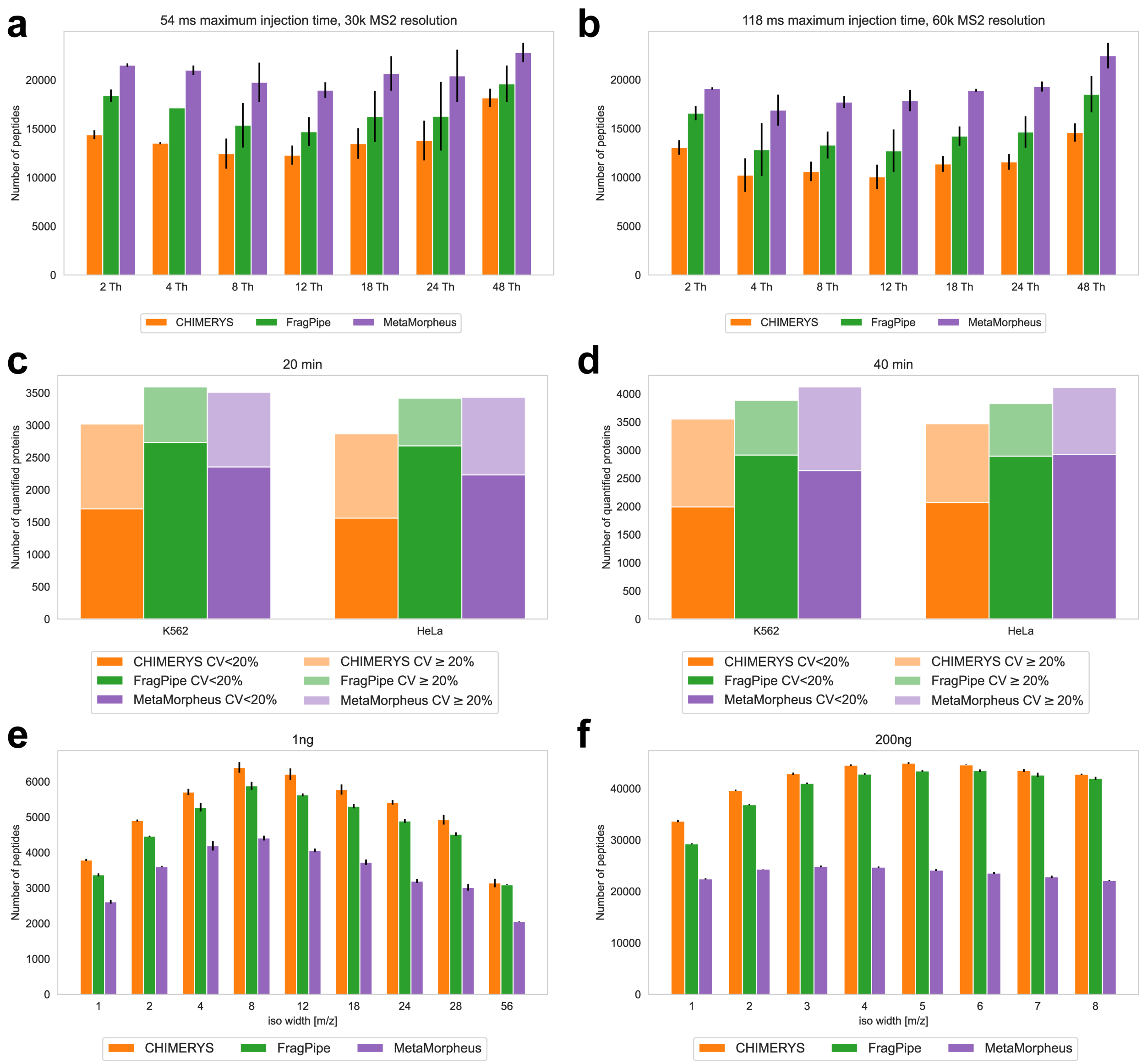


**Supplementary Figure 3. Sensitivity assessment using three WWA datasets. (a)** and **(b)** Numbers of peptides from the first WWA dataset. There are 14 samples with different isolation windows, maximum injection time, and MS2 resolutions. Each sample contain two technical replicates. MBR is enabled. **(c)** and **(d)** Numbers and CVs of quantified proteins from the second dataset. The samples are from K562 and HeLa cell lines, respectively. There are four samples with different combinations of cell line and gradient length. Each sample has eight technical replicates. MBR is enabled. **(e)** and **(f)** Numbers of identified peptides from the third WWA dataset. There are 17 samples with different isolation windows and sample amounts. Each sample has three technical replicates.

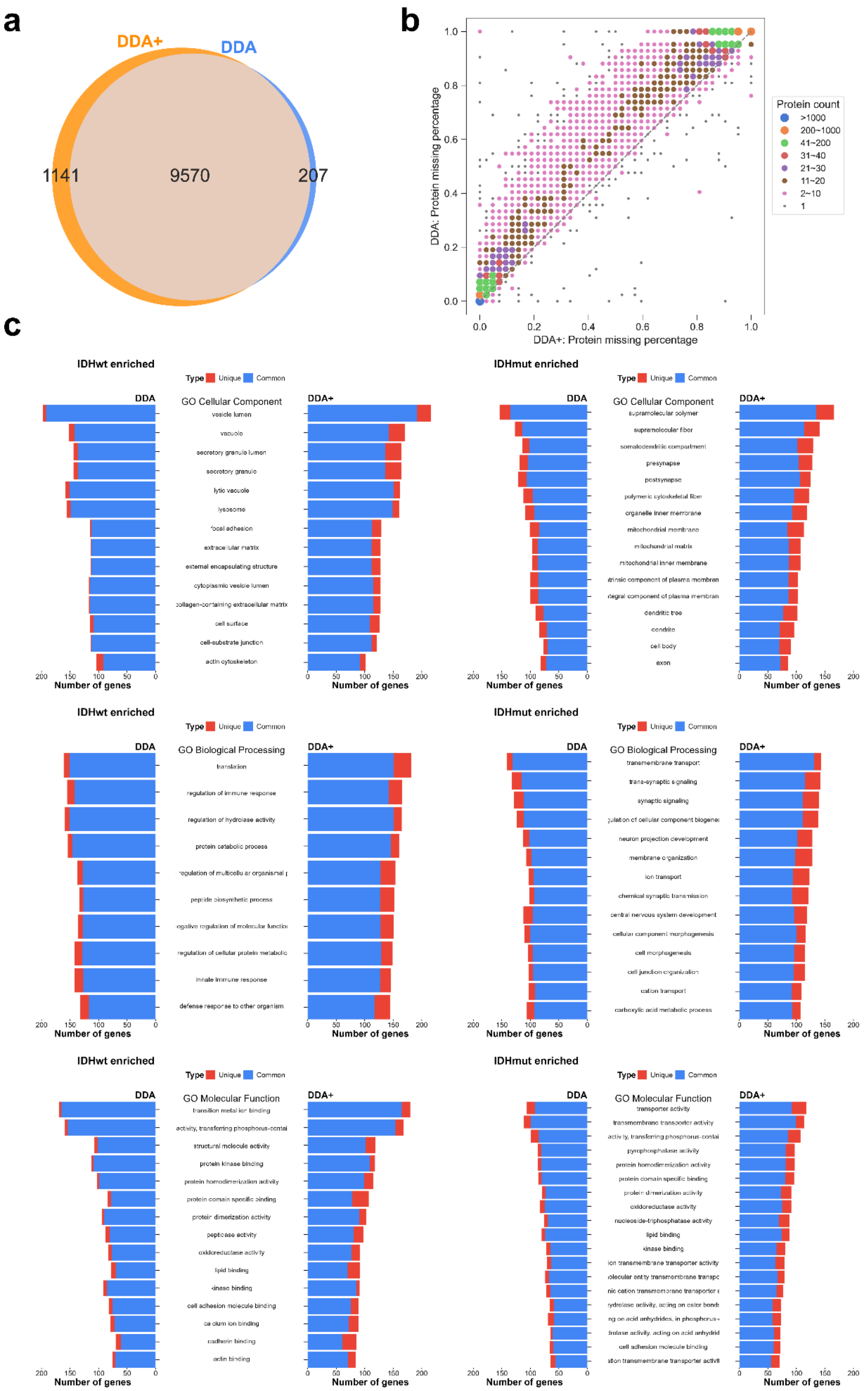
a
DDA+
DDA
1141
9570
207
b
DDA: Protein missing percentage
DDA+: Protein missing percentage
Protein count
>1000
200~1000
41~200
31~40
21~30
11~20
2~10
1
c
IDHwt enriched
IDHmut enriched
Type Unique Common
DDA
DDA+
GO Cellular Component
vesicle lumen
vacuole
secretory granule lumen
secretory granule
lytic vacuole
lysosome
focal adhesion
extracellular matrix
external encapsulating structure
cytoplasmic vesicle lumen
collagen-containing extracellular matrix
cell surface
cell-substrate junction
actin cytoskeleton
supramolecular polymer
supramolecular fiber
somatodendritic compartment
presynapse
postsynapse
polymeric cytoskeletal fiber
organelle inner membrane
mitochondrial membrane
mitochondrial matrix
mitochondrial inner membrane
dendritic tree
dendrite
cell body
axon
GO Biological Processing
translation
regulation of immune response
regulation of hydrolase activity
protein catabolic process
peptide biosynthetic process
innate immune response
defense response to other organism
transmembrane transport
trans-synaptic signaling
synaptic signaling
neuron projection development
membrane organization
ion transport
chemical synaptic transmission
central nervous system development
cellular component morphogenesis
cell morphogenesis
cell junction organization
cation transport
carboxylic acid metabolic process
GO Molecular Function
transition metal ion binding
structural molecule activity
protein kinase binding
protein homodimerization activity
protein domain specific binding
protein dimerization activity
peptidase activity
oxidoreductase activity
lipid binding
kinase binding
cell adhesion molecule binding
calcium ion binding
cadherin binding
actin binding
transporter activity
transmembrane transporter activity
pyrophosphatase activity
nucleoside-triphosphatase activity
ion transmembrane transporter activity
Number of genes

**Supplementary Figure 4. Performance demonstration using a large-scale glioma dataset. (a)** Venn diagram showing the number of quantified genes from the DDA and DDA+ workflows. **(b)** Scatter plot showing the percentage of protein-level missing values by comparing DDA and DDA+ workflows. **(c)** GO analysis using the results from the DDA and DDA+ workflows.
